# AKR1C mediates the acquired resistance to T-Dxd in a HER-2 positive gastric cancer line

**DOI:** 10.1101/2024.12.11.627923

**Authors:** Dong Wang, Qinqin Huang, Peili Wang, Feng Tang, Yuhe Han, Xiangyi Wang, Jingwei Huang, Liqi Shi, Weiqun Cao, Zhixiang Zhang, Qingyang Gu

**Author notes:** Corresponding authors: Zhixiang Zhang.

## Abstract

Trastuzumab deruxtecan (T-Dxd), an anti-HER2 antibody–drug conjugate, has significantly enhanced clinical outcomes for patients as a HER2-directed therapy compared to previous standards of care. However, acquired resistance is always a concern, necessitating further investigation into the underlying resistance mechanism. In this study, we successfully established T-Dxd-resistant cell line (N87-R) by exposing HER2-positive N87 gastric cancer cells to increasing concentrations of T-Dxd, and demonstrated its resistance phenotype both *in vitro* and *in vivo*. While there were no changes in HER2 expression or T-Dxd binding in N87-R, a signature of drug metabolism genes were found upregulated, among which AKR1C played a critical role in the resistance mechanism. The resistance phenotype of N87-R cells was mitigated by both siRNA-mediated knockdown of AKR1C and pharmacological inhibition of its enzymatic activity. Our preclinical study underscores the critical role of AKR1C function in mediating T-Dxd resistance and suggests potential therapeutic innovation for combating T-Dxd resistance.

## Introduction

Gastric cancer ranks among the most prevalent human malignancies globally (1), with approximately 20% of advanced gastric or gastro-oesophageal junction cancers exhibiting HER2 positivity. Research has shown the significance of HER2 overexpression in gastric cancer patients. HER2 expression level correlate with survival of gastric cancer patients (2, 3, 4). Antibody–drug conjugates (ADCs) are a targeted treatment approach that delivers cytotoxic drugs to tumor cells via antibody carriers. Currently, there are 13 FDA-approved ADCs. One of the most notable examples is T-Dxd (DS-8201a), which targets HER2. This ADC includes the anti-human HER2 monoclonal antibody trastuzumab, a cleavable tetrapeptide linker, and the exatecan derivative Dxd (5, 6). T-Dxd is extensively utilized in the treatment of a variety of HER-2 positive cancer types such as breast cancer, gastric cancer, non-small cell lung cancer and has been recently approved for any HER2-positive solid cancer (7, 8, 9). While ADC treatment shows clinical benefit in gastric cancer patients with HER2 expression, primary and secondary ADC resistance to ADC, inevitably emerges (10, 11). Understanding the mechanisms of drug resistance holds promise for refining treatment strategies. Recent studies suggest that multidrug resistance (MDR) regulation (12) or decreased HER2 expression (10) may contribute to T-Dxd resistance, with additional mechanisms potentially contributing to such phenomenon.

We employed N87 cancer cell line, a gastric cancer cell of high HER2 expression, to establish a T-Dxd-resistant clone, which was designated as N87-R. Genomic analysis revealed the upregulation of drug metabolism enzymes in N87-R. We demonstrated the upregulation of AKR1C resulted in reduced anti-tumor effect of T-Dxd, while the genetic knockdown or pharmacological inhibition of AKR1C could restore the drug resistance significantly. Our findings highlighted the importance of AKR1C in mediating the T-Dxd resistance in preclinical setting and suggested the potential of AKR1C inhibition to overcome T-Dxd resistance in future clinical setting.

## Methods

### Reagents

T-Dxd (Trastuzumab deruxtecan, catalog number: HY-138298A), Dxd (catalog number: HY-13631D), DM1 (Mertansine, catalog number: HY-19792), Mefenamic acid (catalog number: HY-B0574), Licochalcone A (catalog number: HY-N0372), Dyclonine hydrochloride (catalog number: HY-B0364A), Exatecan (catalog number: HY-13631) and SN-38 (catalog number: HY-13704) were all bought from MCE. T-DM1 (Trastuzumab Emtansine, catalog number: D4003) was bought from Selleck.

### Mice

Six to eight week old female NOD SCID mice were obtained from Vital River. The mice were maintained in a special pathogen-free environment and in individual ventilation cages (5 mice per cage). All cages, bedding, and water were sterilized before use. The cages with food and water were changed twice a week and all animals had free access to a standard certified commercial laboratory diet. Animal health and behavior were monitored every day.

The experimental endpoint criteria include one of the followings: 1) any animal exhibiting 20% bodyweight loss at any one day 2) Tumor burden exceeds 10% of the animal’s body weight or the mean tumor volume of mouse reaches a value of 3,000 mm^3^ 3) When tumor ulceration occurs, if it does not heal or form a scab within 1 week or is greater than 5 mm diameter or becomes cavitated or if animal develops signs of infection (such as presence of pus) or bleeding, or if the animal shows signs of discomfort (e.g. excessive licking and biting directed at the site) or the animal shows systemic signs of illness (lethargy, decreased activity, decreased food consumption, decreased body condition or weight loss).

All the research staff involved in this animal study have been trained including animal care and procedures. All procedures involving experimental animals were approved by the ethics committee of the WuXi App Tec (Institutional Animal Care and Use Committee Shanghai) and were performed in accordance with the National Guidelines for Animal Usage in Research. Permit number of animal study was ON01-SH006-2023v1.1.

### Cell culture and generation of T-Dxd resistant cell line

N87 cell line (catalog number: CRL-5822) was obtained from the ATCC and cultured in RPMI 1640 medium supplemented with 10% FBS and antibiotics (penicillin 100 U/ml, streptomycin 100 μg/ml) at 37 °C in an atmosphere of 5% CO_2_ in air. Cells were routinely tested for Mycoplasma contamination.

N87 cells were plated in 75 mm plates and treated with 100 ng/ml T-Dxd continuously. Resistant N87-R cell line was obtained 5 months after continuously treatment with T-Dxd *in vitro*. The resistant phenotype was routinely tested by Cell Titer-Glo Luminescent cell viability assay.

### UPLC/MS-MS analysis of Dxd

N87 and N87-R cells were treated with 10 μg/ml Dxd alone or combined with 40 uM mefenamic acid for 24h and 48h. Cells were trypsinized with trypsin and washed with PBS. Then cells were centrifuged and resuspended in PBS to get cell suspension. Cell suspension were precipitated with acetonitrile and further centrifuged. The residue was reconstituted with 150 μL of 10% acetonitrile with 0.1% formic acid and injected into LC-UV-MS for analysis. Experiments were performed on an Orbitrap Eclipse Tribrid mass spectrometer and MS and MS-MS spectra were recorded using Electro Spray Ionization (ESI) in positive ion (PI) and negative ion (NI) mode. Ion transfer tube temperature and vaporizer temperature were set at 325 ℃ and 400 ℃, respectively. Resulting data were analyzed and processed using Xcalibur 4.0 and compound discoverer 3.1 software.

### Cell viability assay

N87 and N87-R cells were seeded on 96-well plates at a density of 20000 cells per well. Cells were then treated with serially diluted concentrations of T-Dxd, Dxd, T-DM1, DM1, Exatecan, SN-38, Mefenamic acid, Licochalcone A, Dyclonine hydrochloride for 7 days and cell viability was detected by Cell Titer-Glo Luminescent Cell Viability Assay Kit (Promega, catalog number: G7573).

### T-Dxd binding assay

2*10^5^ N87 and N87-R cells were incubated on ice for 30 min with different concentrations of T-Dxd. After washing with cold PBS, cells were incubated on ice for 30 min with Goat Anti-Human IgG Fc (DyLight 488, Abcam, ab97003). Then cells were washed with PBS and analyzed by flow cytometry. Data analysis was performed using the FlowJo 10 software.

### Protein expression analysis by flow cytometry

2*10^5^ N87 and N87-R cells were incubated on ice for 30 min with PE anti-human HER2 antibody (Biolegend, clone 24D2, cat#324406), APC anti-human ABCG2 antibody (Biolegend, clone 5D3, cat#332020), PE anti-human ABCB1 Antibody (Biolegend, clone 4E3.16, cat#919406), FITC Mouse Anti-Human MRP1 antibody (BD Biosciences, clone QCRL-3, cat#557593). Then cells were washed with PBS and analyzed by flow cytometry. Data analysis was performed using the FlowJo 10 software.

### Immunohistochemistry

N87 parental cell and N87-R cell subcutaneous tumors were fixed by formalin and paraffin-embedded tissue blocks were sectioned 4 µm thickness. Slides were dewaxed, rehydrated, and blocked for endogenous peroxidase activity. Antigen retrieving was performed in bond epitope retrieval solution (pH 9.0) at 100 °C for 30min. Slides were incubated with peroxide at room temperature for 5 min. Nonspecific antibody binding was blocked by incubating with animal non-immune serum for 10 minutes at room temperature. Slides were then incubated with anti–HER2 (CST, cat#2242, 1:50 dilution) at room temperature for 120 min. Slides were then washed and incubated with biotin-conjugated secondary antibodies for 30 minutes, followed by incubation with avidin DH-biotinylated horseradish peroxidase complex for 30 minutes (Vectastain ABC Elite Kit; Vector Laboratories). The sections were developed with the DAB solution (Leica) and counterstained with hematoxylin. Nuclear staining cells was scored and counted in different vision areas. Images were taken with an OLYMPUS BX43 microscope.

### siRNA analysis

N87-R cells were transfected with two different siRNAs (siAKR1C#1: 5’-GCAUCAGACAGAACGUGCAGGUGUUTT-3’ and siAKR1C#2: 5’-GAGUUCCAGUUGACUGCAGAGGACATT-3’) or NC siRNA to knockdown the expression of AKR1C genes. AKR1C gene expression was evaluated by qPCR 24h post transfection and AKR1C protein expression was evaluated by western blot 48h post transfection. For cell viability assay, AKR1C-silenced cells (24h post transfection) were seeded on 96-well plates at a density of 20000 cells per well. Cells were then treated with serially diluted concentrations of T-Dxd for 6 days and detected by Cell Titer-Glo Luminescent Cell Viability Assay Kit (Promega, catalog number: G7573).

### qPCR analysis

RNA was extracted from N87 or N87-R cells using RNA later reagent, glycogen and sodium acetate. RNA was reverse transcribed to cDNA using M-MLV Reverse Transcriptase. qPCR was performed using SYBR green premix pro Taq HS qPCR kit with a LightCycler 96 instrument (Roche) and PCR was performed using 2×TransTaq-T PCR SuperMix. Relative mRNA levels were calculated using the 2^-ΔΔCt^method and normalized to actin mRNA. Target gene primers are: UTG1A6, 5’-GTATGAAGAACTCGCATCAGC-3’(forward) and 5’-TAGGCTTCAAATTCCTGAGACA -3’ (reverse); ALDH3A1, 5’-AAGATCAGCGAGGCCGTGAA-3’(forward) and 5’-GGCGTTCCATTCATTCTTGTGC-3’ (reverse); AKR1C1, 5’-GGCAATTGAAGCTGGCT-3’ (forward) and 5’-AACTCTGGTCGATGGGAAT-3’ (reverse); AKR1C2, 5’-AGCAAGATTGCAGATGGCAG-3’ (forward) and 5’-CTTCTCCATGGCCTTTACA-3’ (reverse); AKR1C3, 5’-TGGTCCACTTTTCATCGACC-3’ (forward) and 5’-CCAATTGAGCTTTCTTCAGTGAGT-3’ (reverse); GAPDH, 5’-TGGGTGTGA ACCATGAGAAG-3’ (forward) and 5’-GTGTCGCTGTTGAAGTCAGA-3’ (reverse).

### RNA-seq

Total RNA was extracted from the 10^6^ cells using Trizol and Qiagen Rneasy MinElute Cleanup kit (Qiagen) according to the internal SOP instructions in Sequanta Technologies at Shanghai. Then the integrity of the total RNA was determined by 4200 Bioanalyser (Agilent) and quantified using the NanoDrop (Thermo Scientific). About 1000 ng high-quality RNA sample (OD260/280=1.9∼2.0, RIN≥8) was used to construct sequencing library.

RNA purification, reverse transcription, library construction and sequencing were performed in Sequanta Technologies Co., Ltd at Shanghai according to the manufacturer’s instructions. The mRNA-focused sequencing libraries from 1µg total RNA were prepared using TruSeq Stranded mRNA Library Prep kit (Illumina). PolyA mRNA was purified from total RNA using oligo-dT-attached magnetic beads and then fragmented by fragmentation buffer. Taking these short fragments as templates, first strand cDNA was synthesized using reverse transcriptase and random primers. Then in the process of second strand cDNA synthesis, RNA template was removed and a replacement strand, incorporating dUTP in place of dTTP to generate ds cDNA, was synthesized. The incorporation of dUTP quenches the second strand during amplification, because the polymerase does not incorporate past this nucleotide. AMPure XP (Beckmen) beads were then used to purify the ds cDNA from the reaction mix. Next, the ds cDNA was subjected to end-repair, phosphorylation and ’A’ base addition according to Illumina’s library construction protocol. In the following process, Illumina sequencing adapters were added to both size of the ds cDNA fragments. After PCR amplification for DNA enrichment, the AMPure XP Beads were used to clean up the target fragments of 200–300 bp.

After library construction, Qubit 3.0 fluorometer dsDNA HS Assay (Thermo Fisher Scientific) was used to quantify concentration of the resulting sequencing libraries, while the size distribution was analyzed using Agilent BioAnalyzer (Agilent). Sequencing was performed using an Illumina Novaseq 6000 following Illumina-provided protocols for 2×150 paired-end sequencing in Sequanta Technologies.

### Western blot analysis

N87 or N87-R tumor cells were lysed in RIPA Buffer containing 1% protease inhibitor cocktail and 1% phosphatase inhibitor cocktail on ice for 30 min, followed by centrifugation at 12000 *g* for 10 min at 4C. Protein concentrations in the supernatants were measured using BCA protein assay kits, and 50 μg aliquots of proteins were incubated at 95C for 10 min and separated on SDS-PAGE gels. The proteins were transferred to NC (nitrocellulose) membranes, which were blocked in a solution of 5% milk in LI-COR blocking buffer for 1 h at room temperature, and incubated with the indicated primary antibodies overnight at 4C [AKR1C1+AKR1C2+AKR1C3 (ab203834, Abcam, 1:5000 dilution), ALDH3A1 (ab129022, Abcam, 1:2000 dilution), UGT1A6 (NBP2-94532, Novus, 1:500 dilution), Phospho-Chk1 (Ser345) (2348S, CST, 1:1000 dilution), β-Tubulin (86298S, CST, 1:1000 dilution), β-Actin (3700S, CST, 1:1000 dilution)]. The membranes were subsequently incubated with IRDye secondary antibody for 1 h at room temperature, and protein bands were detected by Odyssey CLx imaging system.

### Tumor models and treatment

N87 and N87-R cells (8 million) were mixed with Matrigel (1:1) and injected s.c. into the flanks of 6- to 8-week-old female NOD SCID mice. Tumor-bearing mice (tumor volume around 150 mm^3^) were randomized into 2 groups and treated as below: (1) vehicle control (PBS, i.v.), (2) T-Dxd (3 mg/kg, i.v.) or T-DM1 (5 mg/kg, i.v.). N=6 per group. Tumors were measured using calipers twice a week and tumor volumes were calculated using length*width^2^/2.

N87-R cells (8 million) were mixed with Matrigel (1:1) and injected s.c. into the flanks of 6- to 8-week-old female NOD SCID mice. Tumor-bearing mice (tumor volume around 150 mm^3^) were randomized into 4 groups and treated as below: (1) vehicle control (PBS, i.v.), (2) T-Dxd (3 mg/kg, i.v.), (3) mefenamic acid (25 mg/kg, i.p.) and (4) T-Dxd (3 mg/kg, i.v.)+ mefenamic acid (25 mg/kg, i.p.). N=6 per group. Tumors were measured using calipers twice a week and tumor volumes were calculated using length*width^2^/2. Tumor tissues were harvested after 4 weeks of treatment. The experiment was finished within 5 weeks after the initiation of drug treatment and all animals were humanely euthanized by CO_2_. No animal were found dead before the end of the study.

### Statistical analysis

All statistical analyses were performed using GraphPad Prism 9.0 (GraphPad Software, La Jolla, CA, USA). Results are presented as mean ± SEM and compared by unpaired t-tests, one-way ANOVA or two-way ANOVA. *p* < 0.05 was considered to be statistically significant.

### Data Availability

Correspondence and requests for data underlying the findings or materials should be addressed to Qingyang Gu or Zhixiang Zhang.

## Results

### Generation of T-Dxd-resistant gastric cancer cells

To elucidate the molecular mechanism underlying T-Dxd resistance, we initially generated T-Dxd-resistant clones of the N87 gastric cancer cell line. N87 cells were continuously treated with T-Dxd (13), ultimately yielding T-Dxd-resistant N87 cells (**Fig. 1A**). Subsequently, we evaluated the sensitivity of N87-R cells to Dxd, the payload of T-Dxd. Our findings demonstrated that N87-R cells exhibited resistance to Dxd (**Fig. 1B**). Further investigation involved assessing the response of N87-R cells to T-Dxd treatment in an *in vivo* setting. As presented in **Fig. 1C**, N87-R xenografts displayed resistance to T-Dxd, contrasting with the significant inhibition of tumor weights observed in N87 xenografts upon T-Dxd treatment. Notably, tumor weights of T-Dxd-treated N87-R xenografts did not significantly differ from those of the control-treated group (**Fig. 1D**). Collectively, these results establish the resistance of N87-R cells to T-Dxd both *in vitro* and *in vivo*.

**Fig 1.**
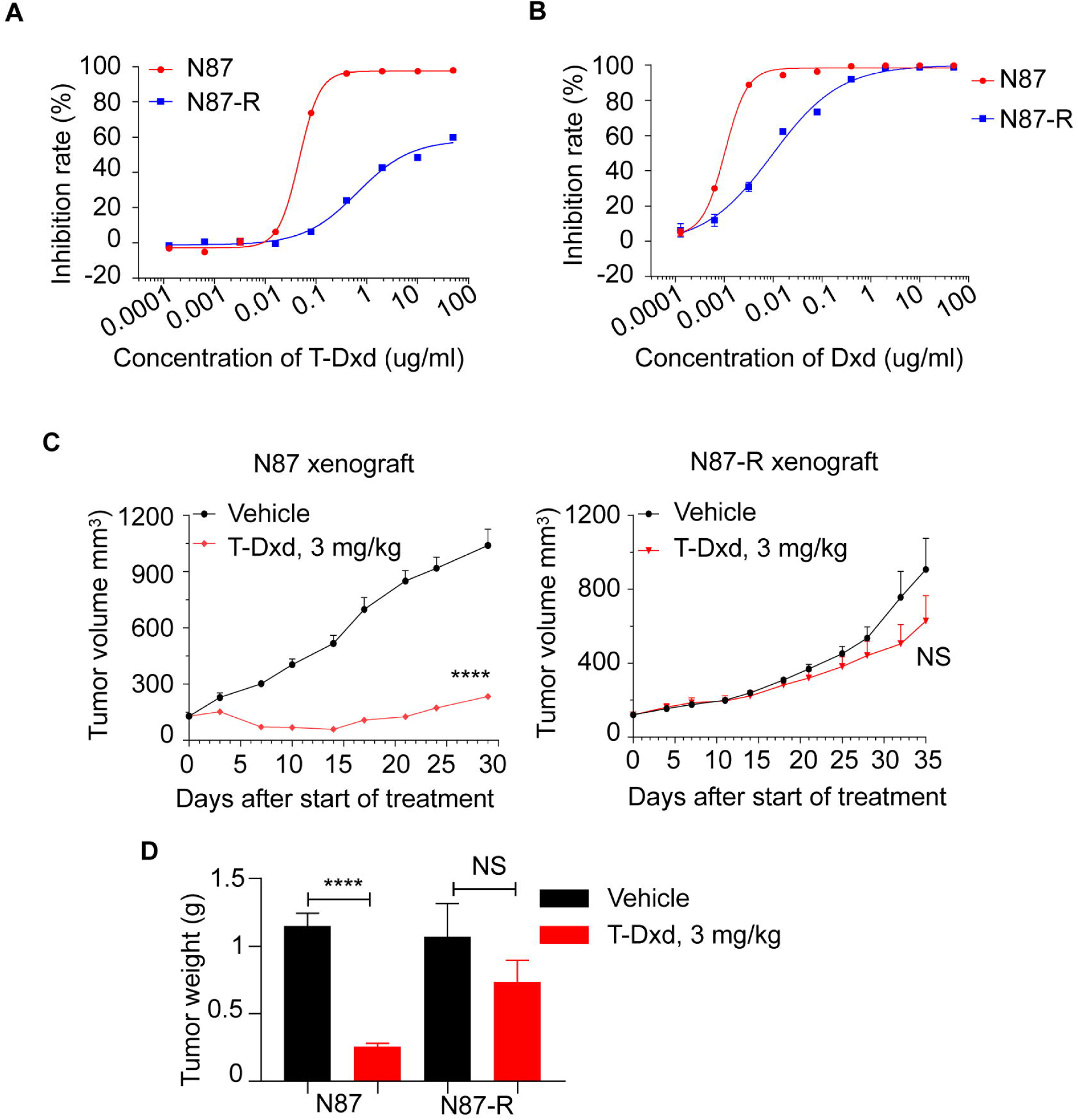
N87-R cells are resistant to T-Dxd *in vitro* and *in vivo*. **(A, B)** N87 and N87-R cells were treated with different concentrations of T-Dxd or Dxd for 7 days. Cell cytotoxicity was detected by CTG. (**C, D**) N87 and N87-R cells were injected subcutaneously to NOD SCID mice and treated with 3 mg/kg T-Dxd or vehicle control. N=6 per group. Tumors were harvested and weighed at the end of the study. Data are shown as the mean ± SEM. Statistical analyses were by two-way ANOVA with Bonferroni comparison test compared with vehicle treated animals (**C**) or by unpaired Student’s t-tests (**D)**. (****, P < 0.0001; NS, P > 0.05). Data are representative from 3 independent experiments.

### HER2 expression, T-Dxd binding and MDR are not involved in T-Dxd resistance in N87-R cells

Numerous potential resistance mechanisms to HER2-directed ADCs have been experimentally investigated, including decreased HER2 expression, reduced ADC binding to tumor cells, and increased expression of drug efflux transporters including ABCG genes (14, 15, 16, 17). To elucidate the resistance mechanism of N87-R cells, we initially assessed HER2 expression in N87 and N87-R cells using flow cytometry. We found no significant difference in HER2 expression between N87 and N87-R cell lines on cell surface level (S1A Fig). Consistent with this result, HER2 expression detected by immunohistochemistry in subcutaneous tumors derived from both N87 and N87-R xenografts also showed no disparity in HER2 expression (S1B Fig). Subsequently, we investigated the binding capability of N87-R cells by incubating them with various concentrations of T-Dxd and evaluating ADC binding to both cell lines. Flow cytometry results indicated no discernible difference in T-Dxd binding between N87 and N87-R cell lines (S1C Fig). Additionally, compared to the parental N87 cell line, N87-R showed no change in expression levels of ABCG1 and ABCG2 (S1D Fig).

To further study the molecular mechanisms involved in T-Dxd resistance, we treated N87 and N87-R cell lines with other HER-2 targeting ADCs and payloads of topoisomerase I inhibitor. CTG cell viability assay showed that both N87 cells and N87-R cells remain sensitive to T-DM1 and DM1 (S1E Fig). Subsequent T-DM1 treatment in N87-R xenografts yielded consistent results with the *in vitro* analysis (S1F Fig). Furthermore, we found that both N87 cells and N87-R cells remain sensitive to other Topoisomerase I inhibitors such as Exatecan and SN38 (S1G Fig). Together, these results indicated that the established N87-R cell line has maintained HER2 and ABCG expression level and T-Dxd binding capability compared to its parental N87 cell line, and developed resistance to T-Dxd via other means.

### Drug metabolism enzymes are upregulated in T-Dxd resistant N87-R cells

To gain further insight into the mechanisms underlying T-Dxd resistance in N87-R cells, we conducted RNA sequencing analysis comparing N87-R cells to their parental N87 counterparts. Our analysis revealed 1124 differentially expressed genes (DEGs) between N87-R and N87 cells (**Fig. 2A**). KEGG analysis of the upregulated DEGs indicated enrichment for pathways associated with xenobiotic metabolism by cytochrome P450 and steroid hormone biosynthesis (**Fig. 2B, 2D**). Focusing on drug metabolism-related genes, both our RNA-seq data and subsequent qPCR analysis demonstrated significantly elevated expression levels of UGT1A6, ALDH3A1, AKR1C1, AKR1C2, and AKR1C3 (21) in N87-R cells compared to N87 cells (**Fig. 2C, 2E**). Furthermore, western blot analysis confirmed these findings, revealing substantially higher protein levels of UGT1A6, ALDH3A1, AKR1C1, AKR1C2, and AKR1C3 in N87-R cells relative to N87 cells (**Fig. 2F**). Collectively, these results suggest that acquired upregulation of drug metabolism signaling occurs in T-Dxd-resistant gastric cancer cells.

**Fig 2.**
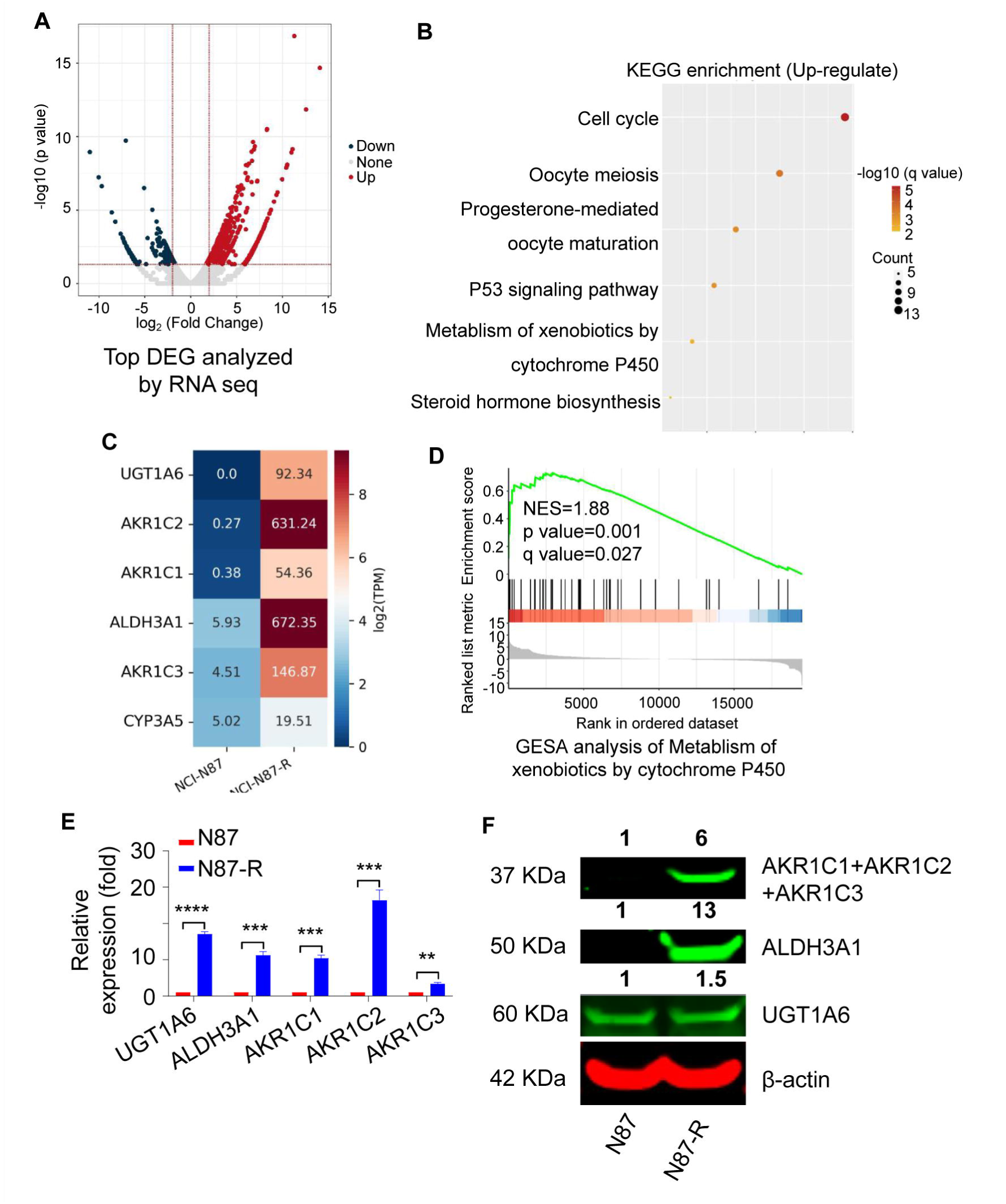
Metabolism of xenobiotics by cytochrome P450 signal is upregulated in T-Dxd resistant N87 cells. **(A)** Volcano plot of genes with differential levels in N87-R cells vs N87 cells. (**B)** KEGG enrichment analysis of upregulation signals in N87-R cells compared with N87 cells. **(C)** Genes with differential levels of metabolism of xenobiotics by cytochrome P450 signal in N87-R cells vs N87 cells. **(D)** GSEA analysis of metabolism of xenobiotics by cytochrome P450 signal. (**E)** qPCR measurements of UGT1A6, ALDH3A1, AKR1C1, AKR1C2, AKR1C3 mRNA levels in N87 and N87-R cells. N=3 per group. Data are shown as the mean ± SEM. Statistical analyses were by unpaired Student’s t-tests. ****, P < 0.0001; ***, P < 0.001; **, P < 0.01. **(F)** UGT1A6, ALDH3A1, AKR1C protein levels in N87 and N87-R cells were examined by western blot. Data are representative from 3 independent experiments (**E, F**).

### AKR1C confers resistance to T-Dxd in gastric cancer cells

To deepen our understanding of the resistance mechanism, we proceeded to investigate the impact of AKR1C on tumor cells. N87-R cells were transiently transfected with either NC (negative control) siRNA or AKR1C siRNA. The efficacy of AKR1C downregulation by siRNA was validated through qPCR and western blot analysis (**Fig. 3A, 3B**). Subsequently, N87-R cells were transfected with NC siRNA or AKR1C siRNA, followed by treatment with T-Dxd for 6 days. While N87-R cells exhibited resistance to T-Dxd, the knockdown of AKR1C expression by siRNA increased the sensitivity of N87-R cells to T-Dxd (**Fig. 3C, 3D**). Taken together, these findings indicate that the downregulation of AKR1C increases sensitivity to T-Dxd treatment in T-Dxd-resistant gastric cancer cells.

**Fig 3.**
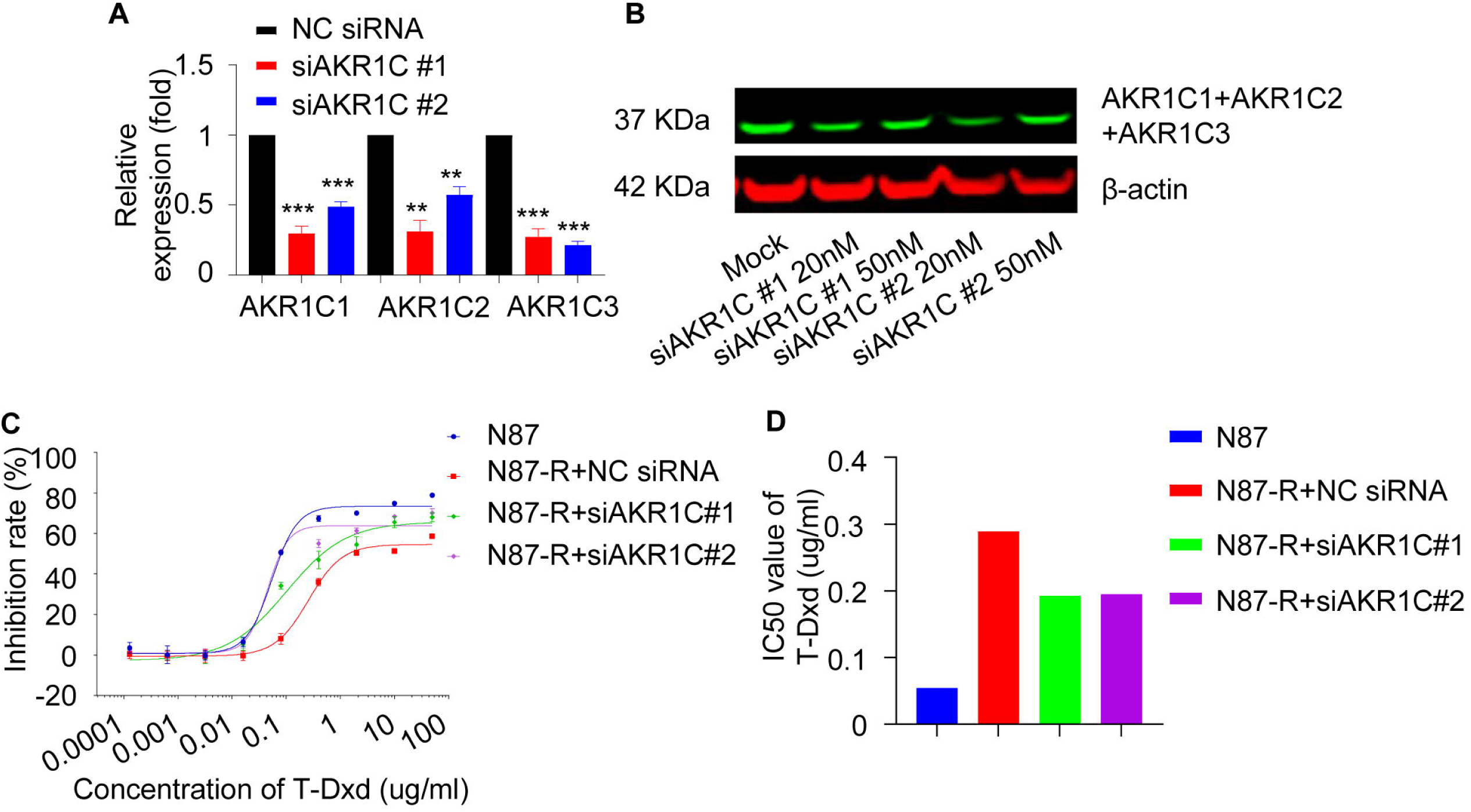
AKR1C confers resistance to T-Dxd in N87 cells. **(A)** N87 or N87-R cells were transfected with NC siRNA or AKR1C siRNA (#1 and #2), cells were collected 24h later. qPCR measurement of AKR1C1, AKR1C2, AKR1C3 mRNA levels was performed. N=3 per group. Data are shown as the mean ± SEM. Statistical analyses were by unpaired Student’s t-tests. ***, P < 0.001; **, P < 0.01. (**B)** N87 or N87-R cells were transfected with NC siRNA or AKR1C siRNA (#1 and #2), cells were collected 48h later. Whole-cell lysates were subjected to AKR1C western blot analysis. (**C, D)** N87 or N87-R cells were transfected with NC siRNA or AKR1C siRNA, cells were collected 48h later following treatment with T-Dxd. Cell cytotoxicity was determined 6 days after T-Dxd treatment. **C**, Cell viability. Data are shown as the mean ± SEM. **D,** IC50 value of T-Dxd. Data are representative from 3 independent experiments.

### Inhibitor of AKR1C overcomes T-Dxd resistance

Mefenamic acid, a nonsteroidal anti-inflammatory drug used for pain and inflammation reduction, has been demonstrated to inhibit AKR1C activity (22, 23). To further elucidate the role of AKR1C in T-Dxd resistance, we initially utilized mefenamic acid to inhibit AKR1C activation and evaluated its effects on the response of gastric cancer cells to T-Dxd treatment *in vitro*. Combination therapy with mefenamic acid and T-Dxd significantly inhibited the growth of N87-R cells and reduced the IC50 value of T-Dxd in N87-R cells compared to T-Dxd monotherapy (**Fig. 4A, 4B**). Additionally, we investigated the impact of combination with other inhibitors, including Licochalcone A, a UGT1A6 inhibitor (24) and Dyclonine hydrochloride, an ALDH3A1 inhibitor (25). However, the combination of Licochalcone A or Dyclonine hydrochloride did not significantly affect the IC50 value of T-Dxd in N87-R cells compared to T-Dxd monotherapy (S2 Fig). To further validate the effect of AKR1C on payload content in N87-R cells, we quantified the Dxd amount in N87 or N87-R cells treated with T-Dxd alone or in combination with mefenamic acid using LC-UV-MS. The results demonstrated that the combination of mefenamic acid with T-Dxd significantly reversed the decrease in Dxd content in N87-R cells (**Fig. 4C**). Prior research has shown that T-Dxd induces the phosphorylation of Chk1, a marker of DNA damage, leading to cell death (4). Subsequent western blot analysis of N87-R cells treated with T-Dxd alone or in combination with mefenamic acid revealed increased DNA damage with combination therapy and mefenamic acid alone did not affect Chk1 phosphorylation (**Fig. 4D**). Furthermore, we evaluated whether the combination of AKR1C inhibitor and T-Dxd could overcome resistance to T-Dxd treatment *in vivo*. Treatment of N87-R xenografts with different drugs demonstrated that while N87-R tumors were resistant to T-Dxd treatment alone, the combination of mefenamic acid and T-Dxd increased their sensitivity (**Fig. 4E**). Collectively, these findings indicate that inhibition of AKR1C by mefenamic acid reverses T-Dxd resistance in gastric tumor cells.

**Fig 4.**
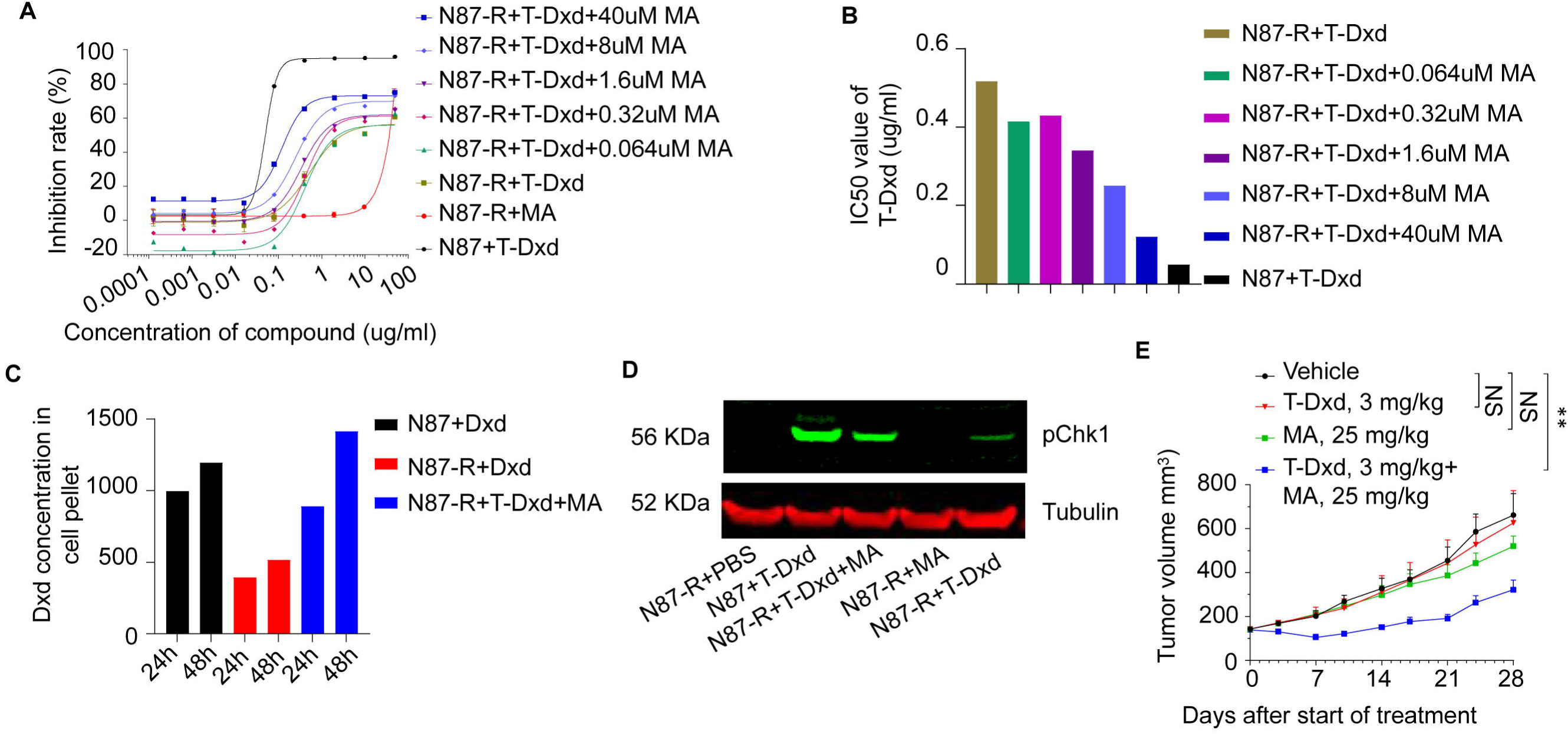
Inhibitor of AKR1C overcomes T-Dxd resistance in gastric tumor cells. (**A, B)** N87-R cells were treated with different concentrations of T-Dxd combined with MA or different concentrations of MA alone for 7 days. **A**, Cell cytotoxicity was detected by CTG. Data are shown as the mean ± SEM. **B,** IC50 value of T-Dxd. (**C)** N87 and N87-R cells were treated with 10 μg/ml Dxd alone or combined with 40 uM mefenamic acid for 24h or 48h, and levels of Dxd in the cell extracts were analyzed by LC/MS. (**D)** N87 and N87-R cells were treated with 10 μg/ml T-Dxd or 40 uM MA for 3 days. Whole-cell lysates were subjected to pCHK1 western blot analysis. (**E)** N87-R cells were injected subcutaneously to NOD SCID mice and treated with 3 mg/kg T-Dxd or vehicle control or 25 mg/kg MA. MA is short for Mefenamic acid. Data are shown as the mean ± SEM. N=6 per group. Tumor volume was assessed by a two-way ANOVA with Bonferroni comparison test compared with vehicle treated animals (**, P < 0.01; NS, P > 0.05). Data are representative from 3 independent experiments.

## Discussion

Human epidermal growth factor receptor 2 (HER2) is a therapeutic target with a long history in anticancer drug development, particularly for breast and gastric cancers. Over 15% of patients with breast cancer, gastric cancer, gastroesophageal junction cancer, lung cancer, biliary tract cancer, pancreatic cancer, prostate cancer, and ovarian cancer exhibit HER2 overexpression (26). The current classification of HER2-targeted drugs includes antibodies, tyrosine kinase inhibitors (TKIs) and antibody-drug conjugates (ADCs) (27). Currently, there are two FDA-approved ADCs, T-DM1 and T-Dxd, for the treatment of HER2-positive cancers. Both of them have demonstrated clinical efficacy in these patient populations. However, some patients develop progressive disease under ADCs due to the occurrence of acquired mechanisms of resistance. The mechanisms of resistance to T-DM1 can be attributed to the antibody or the payload. Mechanisms such as decreased HER2 expression and upregulation of drug efflux pumps like ABCB1 and ABCG2 are known to contribute significantly to resistance to T-DM1 (28, 29). In addition, metabolizing enzymes have also been implicated in mediating chemotherapy resistance (30, 31). Despite few reported mechanisms of T-Dxd resistance, numerous combinational therapy strategies are being evaluated to overcome resistance to anti-HER2 ADCs and improve clinical outcomes in patients, including combinations with tyrosine kinase inhibitors, immune checkpoint inhibitors, or DNA-damaging agents (32, 33, 34). Therefore, gaining a more comprehensive understanding of the molecular mechanisms underlying T-Dxd resistance may facilitate the development of potential therapeutic strategies to overcome this resistance. Thus, we established a T-Dxd resistant model, namely N87-R, using HER2-high-expression gastric cancer N87 cells subjected to chronic exposure to T-Dxd. In our experiment, we demonstrated that AKR1C is overexpressed in N87-R cell line. Furthermore, siRNA-mediated downregulation of AKR1C and inhibition of AKR1C by mefenamic acid sensitize N87-R to T-Dxd treatment, effectively overcome T-Dxd resistance.

Aldo-keto reductases (AKRs) are monomeric NADPH-dependent oxidoreductases that play pivotal roles in the biosynthesis and metabolism of steroids in humans. Specifically, AKR1C enzymes, serving as 3-keto-, 17-keto-, and 20-keto-steroid reductases, are implicated in the pre-receptor regulation of ligands for the androgen, estrogen, and progesterone receptors, as well as in resistance to cancer chemotherapeutic agents such as platin based drugs and topoisomerase inhibitors (35, 36, 37, 38). It has been reported that AKR1C can eliminate irinotecan, which is a topoisomerase I inhibitor, via carbonyl reduction, detoxifying the reactive aldehyde and increasing the production of reactive oxygen species (ROS) (39, 40). AKR1C has been shown to regulate the signaling pathway, increasing AKT phosphorylation, which in turn alleviates reactive oxygen species (ROS) in tumor cells, ultimately leading to chemotherapy resistance (41, 42). More importantly, it is clinically evident that high AKR1C expression correlates with chemotherapy resistance and shorter overall survival in patients with small cell lung cancer or acute lymphoblastic leukemia (43, 44). In line with the previous findings, our result demonstrated that the combination of an AKR1C inhibitor with T-Dxd reversed the decrease in Dxd exposure in N87-R cells, suggesting that AKR1C affected the metabolism process associated with the T-Dxd payload, Dxd, in tumor cells. The discovery of the role of AKR1C in T-Dxd-resistant tumor cells suggested a novel mechanism underlying the development and progression of resistant gastric cancer.

The payload exerts the ADC’s cytotoxic activity and diverse payloads have been used in ADCs including microtubule inhibitors, DNA alkylating reagents and topoisomerase inhibitors (45, 46). To date, two topoisomerase 1 inhibitors have been successfully conjugated to antibodies and approved: the derivative of exatecan, Dxd and the active metabolite of irinotecan, SN-38 (47, 48). In this study, we found the T-Dxd induced resistant N87 cells showed resistance to Dxd but showed no significant resistance to other topoisomerase 1 inhibitors such as exatecan and irinotecan. The difference of structure among Dxd and other topoisomerase 1 inhibitors may lead to the diverse sensitivity of tumor cells. Given the multiple target antigens and expansion of ADC-based therapeutics, our findings of AKR1C upregulation in T-Dxd resistant gastric cancer cells may be relevant to understand resistances raised to other ADCs using Dxd as payload.

Although we provide supporting evidence of the important role of AKR1C to T-Dxd resistance, there are several limitations to this study, and additional considerations which are relevant for clinical translation. The primary setting which we investigated was in one T-Dxd induced resistant gastric tumor cell line. Further study is warranted to determine the generalizability of AKR1C in T-Dxd resistant models such as other T-Dxd resistant gastric cancer cell line or T-Dxd resistant gastric PDX model or even other HER2 positive T-Dxd resistant tumor models.

In summary, our findings indicate that upregulation of AKR1C contributes to resistance to T-Dxd in gastric cancer cells. Inhibition of AKR1C by siRNA or mefenamic acid effectively overcomes resistance to T-Dxd. Moreover, the combined treatment of mefenamic acid and T-Dxd significantly inhibits the growth of T-Dxd-resistant tumors. Our findings suggest that combining AKR1C inhibition may represent a promising treatment strategy for gastric cancer patients resistant to T-Dxd.

## Author’s contributions

D. Wang: Resources, visualization, methodology, writing

Q. Huang: Formal analysis, investigation

P. Wang: Formal analysis, investigation

F. Tang: Investigation, visualization

Y. Han: Investigation, visualization

X. Wang: visualization, writing

J. Huang: Conceptualization

L. Shi: Formal analysis, investigation

W. Cao: Resources, methodology

Z. Zhang: Conceptualization, resources, project administration, writing–review and editing

Q. Gu: Conceptualization, resources, project administration

