## supplementary figure 1 for "AKR1C mediates the acquired resistance to T-Dxd in a HER-2 positive gastric cancer line"

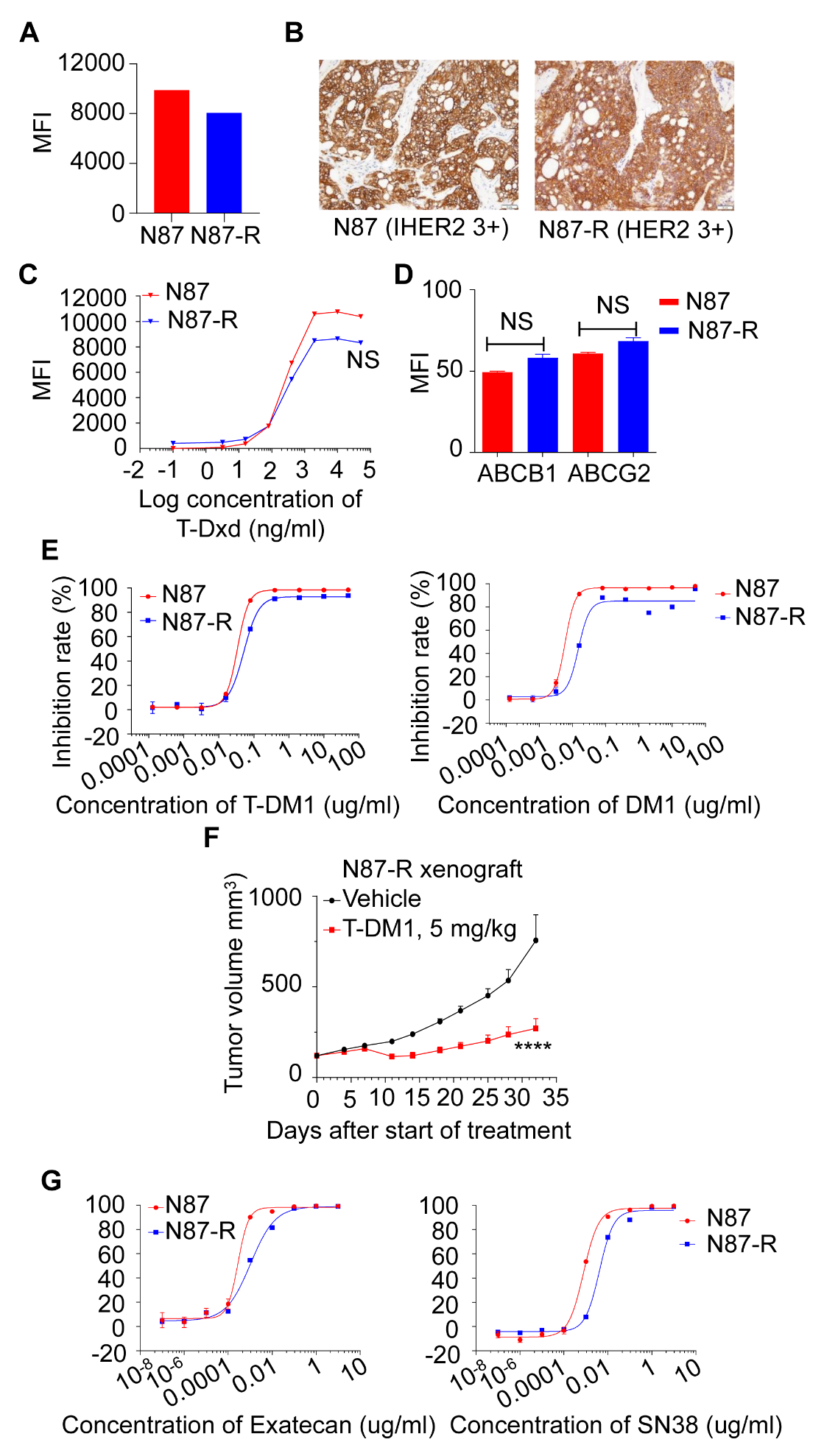

**S1 Fig. HER2 expression, binding, sensitivity of other HER2 targeted ADC and topoisomerase I inhibitors are not significantly different between N87 and N87-R cells.**

**(A)** HER2 expression of N87 and N87-R cells was detected by flow cytometry. **(B)** N87 and N87-R tumor cells were inoculated subcutaneously to mice. Tumors were collected and HER2 protein expression was detected by IHC. Scale bar=50 μm. (**C)** N87 and N87-R cells were treated with different concentrations of T-Dxd on ice for 0.5 h and binding of T-Dxd to cells was analyzed by flow cytometry. Data are shown as the mean ± SEM. Statistical analyses were by two-way ANOVA with Bonferroni comparison test. NS, P > 0.05. (**D)** Expression of multidrug resistance proteins were detected by flow cytometry. Statistical analyses were by unpaired Student’s t-tests. NS, P > 0.05. (**E**) N87 and N87-R cells were treated with different concentrations of T-DM1, DM1 for 7 days. Cell cytotoxicity was detected by CTG. Data are shown as the mean ± SEM. **(F)** N87-R cells were injected subcutaneously to NOD SCID mice and treated with 5 mg/kg T-DM1 or vehicle control. Tumors were harvested at the end of the study. N=6 per group. Data are shown as the mean ± SEM. Statistical analyses were by two-way ANOVA with Bonferroni comparison test. ****, P < 0.0001. **(G)** N87 and N87-R cells were treated with different concentrations of Exatecan, SN38 for 7 days. Cell cytotoxicity was detected by CTG. Data are shown as the mean ± SEM. Data are representative from 3 independent experiments.
